# Neuronal Gene Architecture in *Cancer borealis* Revealed by Long-Read Genome Assembly and Deep Transcriptomic Analysis

**DOI:** 10.64898/2026.08.12.744261

**Authors:** Murugesan Raju, Adam J. Northcutt, David J. Schulz

**Affiliations:** Division of Biological Sciences, University of Missouri, Columbia, MO, USA, 65211

**Keywords:** Neurogenomics, Transcriptome, Genome Annotation, Crustacean

## Abstract

Understanding the underlying neuronal function in non-model organisms requires accurate resolution of gene structure and transcript diversity. Here, we present a comprehensive genome annotation tor the Jonah crab (*Cancer borealis)*, a key experimental system in crustacean neurobiology, with a particular focus on transcriptome-supported neuronal gene architecture. By integrating long-read genome assembly with extensive transcriptomic evidence, we reconstructed gene models with high confidence, enabling detailed characterization of exon-intron organization, alternative splicing, and isotorm diversity across gene families. Functional classification revealed extensive representation of neural-associated gene classes, including ion channels and receptors, transporters, enzymes, zinc finger proteins, histones, structural proteins, and cell adhesion molecules, alongside a large set of previously uncharacterized genes. In this study we particularly focused on the neuronal and ion channel gene families known to underlie circuit-level neuronal function in *C. borealis*. We provide an in-depth analysis of 87 genes spanning 17 neural-related gene families and 41 neuropeptides, detailing chromosomal localization, gene length, exon-intron configuration, and transcript-supported isotorm structure. For many of these genes, transcriptomic data confirmed expression and refined coding boundaries. Comparisons with existing transcriptomic datasets demonstrate strong concordance in gene expression patterns while also revealing novel transcripts and expanded gene family members not previously annotated. Together, this genome and transcriptome-integrated annotation establishes a high-resolution framework tor studying neuronal gene organization in *C. borealis*. T his resource enables direct connections between gene architecture, transcript diversity, and neural function, supporting future investigations in crustacean neurogenomics, comparative genomics, and the evolution of nervous system complexity.

## Introduction

The Jonah crab, *Cancer borealis*, is a brachyuran crustacean distributed along the Atlantic coast of North America from Nova Scotia, Canada, to Florida, USA, occupying habitats from the intertidal zone to depths approaching 800 meters Stehlik *et al*. (1991). Beyond its ecological relevance, *C. borealis* has emerged as a premier experimental system in neurobiology, particularly for elucidating the principles governing neuronal excitability, circuit dynamics, and neuromodulation. As a member of the Decapoda (Arthropoda; Malacostraca), it occupies a key phylogenetic position for comparative studies aimed at understanding the evolution of nervous systems across metazoans Harzsch (2006).

Crustacean model systems, including *C. borealis*, have played a foundational role in modern neuroscience. Seminal work in decapod crustaceans established core concepts such as command neurons and descending control of behavior Wiersma and Ikeda (1964), electrical synapses and their functional roles Furshpan and Potter (1959), and presynaptic inhibition as a mechanism for synaptic modulation Dudel and Kuffler (1961). Subsequent studies in related taxa, including crayfish and lobsters, were instrumental in identifying *γ*-aminobutyric acid (GABA) as a major inhibitory neurotransmitter Otsuka *et al*. (1966, 1967) and in characterizing transient potassium currents (I_A_) that shape neuronal firing and spike frequency adaptation Connor (1975); Connor *et al*. (1977). Additional experimental innovations, such as intracellular dye filling for neuron identification Remler *et al*. (1968); Stretton and Kravitz (1968) and detailed analyses of rhythmic motor circuits Hartline (1967); Maynard (1972); Mulloney and Selverston (1974); Heitler (1978); Marder and Bucher (2007), further established crustaceans as indispensable systems for linking cellular properties to circuit-level function.

Central to these discoveries is the role of ion channels and asso ciated membrane proteins, which collectively determine neuronal excitability and information processing. The selective expression of ion channels across neuron types defines intrinsic electrical prop erties and constrains how neurons respond to synaptic input and neuromodulatory signals Kick and Schulz (2022); Marder (2012); Marder and Taylor (2011). Variation in channel composition, den sity, and kinetic properties enables functional diversity even among morphologically similar neurons, a principle extensively demon strated in crustacean central pattern generator circuits Goaillard and Marder (2009). Thus, understanding the genomic organization and transcriptomic diversity of ion channel genes is essential for linking molecular mechanisms to neural circuit function and behavioral output Schulz *et al*. (2006); O’Leary *et al*. (2013).

Despite their outsized contributions to neuroscience, crustaceans remain underrepresented in genomic databases relative to insects and vertebrates. Early surveys revealed a pronounced imbalance in publicly available molecular data, with orders-of-magnitude fewer Sequence Read Archive (SRA) and Gene Expression Omnibus (GEO) entries for crustaceans than for insects Northcutt *et al*. (2016). Al-though this disparity has narrowed modestly with recent sequencing efforts, comprehensive, chromosome-scale genome assemblies for large-bodied decapods remain scarce. The first published crustacean genome, that of the microcrustacean *Daphnia pulex*, appeared in 2011 Colbourne *et al*. (2011), more than a decade after the completion of the *Drosophila melanogaster* genome Adams *et al*. (2000). This gap has limited the ability to investigate gene architecture, regulatory complexity, and genome evolution in crustacean nervous systems.

Transcriptomic studies of *C. borealis* and related species have begun to address this limitation by identifying expressed genes involved in neuronal signaling, including ion channels, neurotransmitter receptors, and synaptic proteins Schulz *et al*. (2007); Northcutt *et al*. (2016). However, transcriptome-based analyses alone provide an incomplete picture, as they lack information on genomic context, exon–intron organization, long-range gene structure, and chromosomal localization. These features are particularly important for neuronal genes, which often span large genomic regions, contain long introns, and exhibit extensive alternative splicing that contributes to functional diversity Grabowski (2011).

The recent availability of a high-quality, long-read *de novo* genome assembly for *C. borealis* (isolate JC504) represents a major advance for the field Polinski *et al*. (2025). This assembly, generated using PacBio long-read sequencing in combination with Omni-C scaffolding, achieves chromosome-level resolution, spanning 691 Mb and organized into 51 chromosome-length scaffolds with a limited number of unplaced contigs. Such contiguity enables precise mapping of genes, resolution of complex loci, and integration of transcriptomic evidence to refine gene models.

High-resolution genome assemblies are transformative for studies of nervous system biology, as they enable analyses of gene architecture that are inaccessible through transcriptomics alone. Gene length, intron size, exon number, and alternative splice site usage can influence transcriptional regulation, RNA processing, and protein diversity Pandya-Jones and Black (2009). In neurons, where functional specialization often depends on finely tuned expression of multiple isoforms, these architectural features are critical for understanding how genomic organization contributes to physiological diversity and robustness Keren *et al*. (2010).

In this study, we leverage the chromosome-scale genome assembly of *C. borealis* together with extensive transcriptomic evidence to generate a refined, genome-anchored annotation with a specific focus on neuronal genes. Building on previously identified neural gene sets Northcutt *et al*. (2016), we characterize exon–intron organization, gene span, isoform diversity, and chromosomal distribution for key neuronal and ion channel gene families. We then perform a de novo RNAseq analysis from pooled stomatogastric ganglia to validate the utility of this enhanced assembly relative to an existing transcriptome assembly Northcutt *et al*. (2016) for quantitative analysis. By doing so, we generate novel data towards the potential importance of previously unstudied ion channel and receptor proteins via their relative abundance in these ganglia. By anchoring transcriptomic findings within a genomic framework, this work establishes a molecular foundation for future investigations into crustacean neurobiology, comparative neurogenomics, and the evolutionary principles governing neuronal gene architecture.

## Materials and Methods

### Tissue Collection and RNA Preparation

Adult male crabs, *Cancer borealis*, were obtained from The Fresh Lobster Company (Gloucester, Massachusetts, USA) and maintained in artificial seawater at 12°C until used. Crabs were anesthetized by packing them in ice for 30 minutes. The complete stomatogastric nervous systems (STNS) (including the commissural, esophageal, and stomatogastric ganglia) were dissected and pinned out in a Sylgard (Dow Corning)–coated dish containing chilled (12–13 °C) physiological saline. These were then further manually cleaned of connective tissue and muscle to the extent possible, and the tissues were rinsed several times in physiological saline made with ultrapure, RNase free water. After dissection, whole stomatogastric ganglia from 15 different individuals were pooled and homogenized in Trizol (Invitrogen). Insoluble tissues were pelleted by centrifugation, the supernatant removed and RNA extraction was performed as per the protocol provided by the manufacturer (Invitrogen), and subsequently treated with DNase (Zymo Research) prior to library construction.

### Stomatogastric Ganglion (STG) RNAseq and de novo Transcriptome Assembly

Library construction and RNA-sequencing were performed by the Genomics Technology Core (RRID:SCR017778) at the University of Missouri (Columbia, MO, USA). Briefly, RNA samples were quantified using Qubit 2.0 Fluorometer (Life Technologies, Carlsbad, California, USA) and RNA integrity checked using an Agilent 2100 Bioanalyzer (Agilent Technologies, Palo Alto, California, USA). mRNAs were purified using poly-T oligo-attached magnetic beads and then fragmented. The first and the second strand cDNAs were synthesized and end repaired. Adaptors were ligated after adenylation at the 3’ends. Then cDNA templates were enriched by PCR. cDNA libraries were validated using a High Sensitivity Chip on the Agilent 2100 Bioanalyzer. The cDNA library was quantified using Qubit 2.0 Fluorometer (Life Technologies, Carlsbad, California, USA). The samples were then loaded on the Illumina HiSeq 2000 instrument for sequencing with a 2×100 paired-end configuration

### Genome Assemblies and Merging Strategy

We performed secondary analysis of two independently generated genome assemblies for *Cancer borealis* were used to produce a unified reference genome: GCA_041682235.1 (GMGI Cborealis 1.0; BioProject PRJNA1097536) generated by Polinski *et al*. (2025) and GCA_036785275.1 (qmCanBore1 p1.0; BioProject PRJNA1018997) generated by the Canadian BioGenome Project https://www.ncbi.nlm.nih.gov/datasets/genome/?bioproject=PRJNA1018997. These assemblies were available in NCBI Genomes. To assess concordance and identify syntenic regions, whole-genome alignments were performed using NUCmer from the MUMmer v4.0.0 package Marçais *et al*. (2018). Alignments were generated with parameters optimized for large, repeat-rich metazoan genomes. Resulting .delta files were processed with show-coords to identify collinear blocks and structural discrepancies.

Custom scripts were used to resolve conflicts and generate a merged assembly by prioritizing contiguity, structural consistency, and alignment support. Assembly statistics of the merged genome closely matched those of GMGI Cborealis 1.0, while incorporating additional resolved regions from the complementary assembly. The merged genome served as the reference for all downstream analyses.

### RNA-Seq Data Processing and Alignment

The RNA-seq datasets derived from multiple *C. borealis* tissues, including stomatogastric ganglion (STG) neurons, were used to provide transcriptomic evidence. Raw reads quality were assessed using FastQC v0.12.0 Andrews (2010). Adapter sequences and low-quality bases were removed using *Trimmomatic* (v0.401) (Bolger *et al*. 2014)

### Gene Prediction Using RNA-Seq Evidence

Gene models were predicted using BRAKER2 v2.1.6 Brna *et al*. (2021), which integrates *ab initio* prediction with RNA-seq-derived intron evidence. The soft-masked genome assembly and RNA-seq BAM files were used as input. GeneMark-ET provided self-training models based on detected introns, which were subsequently refined using AUGUSTUS with RNA-seq hints. This approach improved exon–intron boundary accuracy and supported alternative isoform detection. Final gene predictions were exported in GTF format.

### Prediction of Protein-Coding Regions

Candidate protein-coding sequences were identified using TransDecoder v5.5.0 Haas *et al*. (2013). Transcript sequences were extracted from the BRAKER2 annotation using `gffread`. Long open reading frames were identified based on length and codon usage statistics, followed by refinement using homology support from BLASTp searches against the NCBI non-redundant (nr) protein database and Pfam domain identification Finn *et al*. (2016). Only high-confidence ORFs with coding potential or conserved domains were retained.

### Functional Annotation

Predicted protein sequences were functionally annotated using DI-AMOND BLASTp v2.1.6 Buchfink *et al*. (2021) against the NCBI nr database. Annotations were assigned based on best hits, requiring an e-value < 1 × 10−5, minimum alignment length of 50 amino acids, and percent identity greater than 30%.

### Genome Completeness Assessment

Annotation completeness was assessed using BUSCO v5.3.2 Manni *et al*. (2021) with the *arthropoda_odb10* dataset. Complete, duplicated, fragmented, and missing orthologs were quantified to evaluate genome and annotation quality.

### EGAPx-Based Genome Annotation

To further refine gene models, we employed EGAPx, the public implementation of the NCBI Eukaryotic Genome Annotation Pipeline Goldfarb *et al*. (2024). The polished genome assembly and taxonomic identifier for *C. borealis* (taxid: 39395) were provided as input. EGAPx integrates protein homology alignments via miniprot, RNA-seq alignments via STAR Dobin *et al*. (2013), and *ab initio* predictions using species-appropriate hidden Markov models. These evidence sources were integrated by Gnomon to generate final gene models, which were output as GFF and GTF files.

### Genome-Anchored Neuronal Gene Annotation

A curated list of neuronal genes previously identified from deep transcriptomic profiling Northcutt *et al*. (2016) served as the foundation for focused analysis. These transcripts were mapped to the chromosome-scale genome assembly Polinski *et al*. (2025) to define genomic boundaries, exon–intron structure, isoform organization, and chromosomal location.

### Extraction and Analysis of Gene Architecture Features

Key genomic features were extracted for each neuronal gene, including exon count, total exon length, genomic span, and chromosomal coordinates. Genomic span was calculated as the distance between gene start and end positions, encompassing intronic and regulatory regions. Custom Python scripts parsed GTF files and calculated summary statistics using pandas v2.2.3. Data visualization was performed using seaborn v0.13.2 and matplotlib v3.10.3..

### Transcriptome to Genome Alignment

To evaluate the quality and structural completeness of the *de novo* transcriptome assembly, the assembled transcripts were mapped against the *Cancer borealis* reference genome. Spliced sequence alignment was performed using Minimap2 (version 2.28-r1209) utilizing the `-ax splice` parameter, which is specifically designed to accommodate transcript-to-genome mapping by allowing alignments to span intronic regions. The resulting alignment file was processed using SAMtools to convert, sort by genomic coordinates, and index the data. Finally, primary and overall mapping statistics were calculated using the `samtools flagstat` utility to quantify the proportion of the transcriptome represented in the reference genome.

### Transcriptome Quantification Methods

To generate robust expression estimates and comprehensively evaluate the STG transcriptome dataset, we applied three complementary quantification strategies: a *de novo* assembly and quantification workflow using Trinity and RSEM, a reference-guided genome-mapping approach using STAR and featureCounts, and an independent proprietary *de novo* assembly and quantification workflow using DNASTAR.

#### De novo Transcriptome Assembly and Expression Quantification Using Trinity

Raw sequencing reads in FASTQ format were subjected to quality control and adapter trimming to remove low-quality bases and sequencing artifacts using *Trimmomatic* (v0.401) (Bolger *et al*. 2014). Prior to assembly, contaminating sequences originating from bacterial, viral, and other non-target taxa were identified and removed by sequence-similarity filtering against public databases. The resulting high-quality, filtered reads were then used for *de novo* transcriptome assembly with Trinity (v2.15.2) (Grabherr *et al*. 2011; Haas *et al*. 2013) using default parameters. Trinity reconstructs transcript sequences through the sequential operation of its Inchworm, Chrysalis, and Butterfly modules. This *de novo* assembly provided genome-independent transcript evidence for validating predicted gene structures and facilitated the recovery of transcripts that may be fragmented or incompletely represented in the genome assembly.

To assign biological context and functional labels, Trinity-assembled contigs were annotated against the DIAMOND Buchfink *et al*. (2021) to identify putative orthologous groups and functional annotations. In addition, predicted coding sequences were compared with the genome annotation using sequence-similarity searches and reciprocal best-hit analyses to identify corresponding neuronal genes and candidate ion channel transcripts based on Northcutt *et al*. (2016) published article.

For expression quantification, quality-filtered reads were mapped back to the Trinity-assembled transcriptome reference. Transcript abundances were estimated using RSEM (v1.3.3) (Li and Dewey 2011) through the Trinity abundance-estimation workflow. To reduce isoform-level mapping ambiguity and support comparisons across complex neural gene families, transcript-level estimates were summarized at the gene level. Final expression profiles were normalized and reported as transcripts per million (TPM).

#### Reference-Guided Mapping and Quantification Using STAR

To evaluate transcriptional support for the genome annotation, the STG RNA-seq datasets were analyzed using a reference-guided mapping approach. Raw paired-end Illumina reads were subjected to quality control and adapter trimming using *Trimmomatic* (v0.401) (Bolger *et al*. 2014). Quality-filtered reads were aligned to the *Cancer borealis* genome assembly using STAR v2.7 (Dobin *et al*. 2013) in two-pass mode to improve splice-junction detection.

Genome indices were generated from the final genome assembly and the corresponding gene annotation file. During the first alignment pass, splice junctions identified across samples were collected and incorporated into the second-pass alignment strategy. Reads were then realigned using the updated splice-junction information to improve mapping accuracy across exon–intron boundaries.

Gene-level read counts were generated using `featureCounts` from the Subread package v2.0 (Liao et al. 2013). Only uniquely mapped reads were retained for counting. Read assignments were performed against annotated exon features, and counts from exons belonging to the same gene model were summarized to generate gene-level count estimates. The resulting count matrix was used to assess transcriptional support for predicted genes and to compare expression patterns with previously published transcriptomic resources.

#### Proprietary de novo Assembly and Quantification Using DNASTAR

As a third comparative approach, *de novo* assembly and expression quantification were performed using the commercial DNASTAR Lasergene software suite (DNASTAR, Inc., Madison, WI). Raw sequencing reads were preprocessed to remove adapters and low-quality sequences and were then assembled *de novo* using the SeqMan NGen application. Assemblies were generated using default transcriptome parameters optimized for Illumina RNA-seq data, which cluster reads into contigs using proprietary overlap-based algorithms.

Following consensus contig generation, transcript abundance was quantified using the QSeq module within ArrayStar. Preprocessed reads were mapped back to the assembled STG contigs. To enable direct comparison with the Trinity and STAR pipelines, read counts were summarized for each contig and normalized as transcripts per million (TPM).

### Transcriptome Integration and Comparative Expression Analysis

To evaluate the consistency of transcript abundance estimates and provide independent validation of the genome annotation, neuronal gene expression profiles derived from three transcriptomic workflows were compared. Transcript-level abundances were subsequently collapsed to gene-level estimates to facilitate direct comparison with genome-based quantification and previously published transcriptomic data. Gene-level expression values obtained from the DNASTAR, Trinity, and STAR/featureCounts workflows were log_2_(count + 1) transformed prior to comparison. Pairwise concordance among datasets was evaluated using Pearson correlation analysis, and scatter plots were generated to visualize relationships between expression estimates derived from the three independent quantification approaches. This comparative framework provided an assessment of transcriptomic reproducibility and supported the accuracy of neuronal gene annotations identified in the genome assembly.

## Results and Discussion

The integration of deep transcriptomic sequencing Northcutt *et al*. (2016) with a chromosome-scale long-read genome assembly Polinski *et al*. (2025) enables a comprehensive genomic characterization of neuronal genes in the Jonah crab, *Cancer borealis*. Moving from a transcriptome-centric perspective to a genome-anchored framework provides unprecedented resolution of neuronal gene architecture, including exon–intron organization, chromosomal localization, gene span, and paralogous gene relationships. This genome-based view reveals structural and evolutionary features of neuronal genes that are not accessible through transcriptome data alone.

### Genome Assembly Quality and Genomic Context

All analyses were conducted using a high-quality *de novo* genome assembly of *Cancer borealis*, spanning 691.2 Mb and consisting of 157 contigs, with an N50 of 17.18 Mb and a maximum contig length of 46.07 Mb (Table 1). The assembly exhibited a GC content of 40.77%, consistent with values reported for other decapod crustaceans and indicative of a high-quality genomic resource suitable for gene prediction, annotation, and transcriptomic analyses.

**Table 1.**
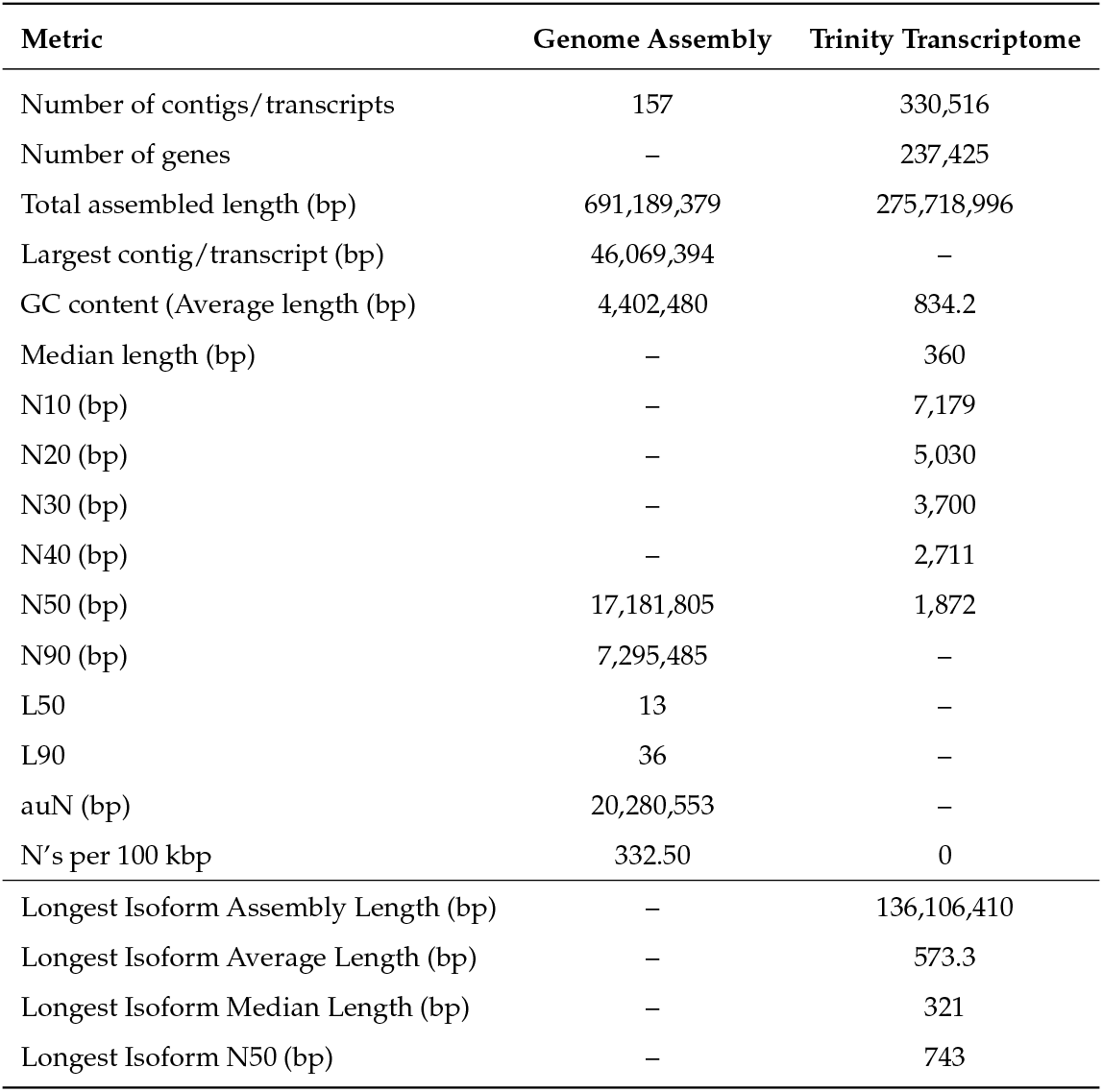
Summary statistics of the *Cancer borealis* genome assembly and Trinity de novo transcriptome assembly.

| Metric | Genome Assembly | Trinity Transcriptome |
| --- | --- | --- |
| Number of contigs/transcripts | 157 | 330,516 |
| Number of genes | – | 237,425 |
| Total assembled length (bp) | 691,189,379 | 275,718,996 |
| Largest contig/transcript (bp) | 46,069,394 | – |
| GC content (Average length (bp)) | 4,402,480 | 834.2 |
| Median length (bp) | – | 360 |
| N10 (bp) | – | 7,179 |
| N20 (bp) | – | 5,030 |
| N30 (bp) | – | 3,700 |
| N40 (bp) | – | 2,711 |
| N50 (bp) | 17,181,805 | 1,872 |
| N90 (bp) | 7,295,485 | – |
| L50 | 13 | – |
| L90 | 36 | – |
| auN (bp) | 20,280,553 | – |
| N's per 100 kbp | 332.50 | 0 |
| Longest Isoform Assembly Length (bp) | – | 136,106,410 |
| Longest Isoform Average Length (bp) | – | 573.3 |
| Longest Isoform Median Length (bp) | – | 321 |
| Longest Isoform N50 (bp) | – | 743 |

To generate independent transcriptional evidence in support of the genome annotation, a de novo transcriptome was assembled from contamination-filtered STG RNA-seq reads using Trinity. The resulting assembly contained 237,425 putative genes and 330,516 transcript isoforms, encompassing 275.7 Mb of assembled sequence with a GC content of 44.39% and a transcript N50 of 1,872 bp (Table 1). Collectively, the genome and transcriptome assemblies constitute complementary resources for the validation of gene models, the identification of neuronal genes and ion channel gene families, and the quantification of transcript abundance across multiple independent analytical workflows.

Genome-wide evaluation metrics, including Nx plots, cumulative contig length distributions, and GC content profiles (Figure 1), demonstrate balanced base composition and high contiguity. These features provide a robust foundation for resolving large neuronal genes with extensive intronic regions and complex exon structures.

**Figure 1.**
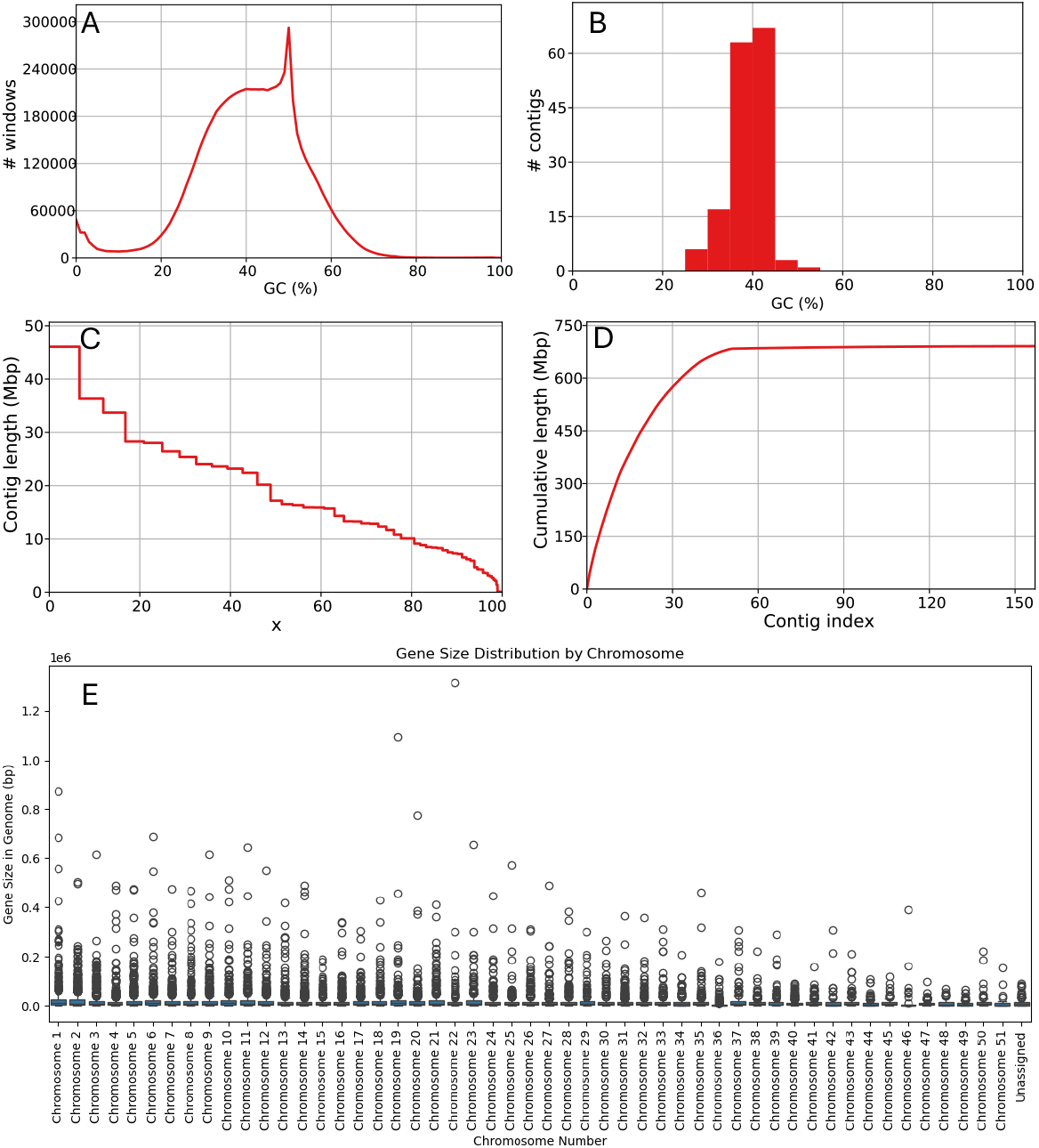
Genome assembly characteristics of *Cancer borealis*. **(A)** Distribution of genomic windows according to GC content. **(B)** Distribution of GC content among assembled contigs. (**C**) Nx curve showing contig length as a function of the percentage of the total assembly represented. (**D**) Cumulative assembly length after ordering contigs from longest to shortest, reaching a total assembly size of approximately 691.2 Mb. (**E**) Distribution of annotated gene lengths across the 51 chromosome-level scaffolds and unassigned contigs; each point represents an individual gene.

### Genome Annotation and Global Gene Architecture

Annotation using the EGAPx workflow identified 17,055 genes and 21,251 transcripts, of which 91.7% are protein-coding. Long non-coding RNAs and other non-coding biotypes account for the remaining 8.3%. Across the genome, exon counts range from one to 210 per gene, with a median of six exons. Mean exon length is approximately 2.16 kb and maximum gene size is extending beyond 1.3 Mb(Table 2 Gene Annotation Metrics(A)).

**Table 2.** Overview of gene annotation and functional categories in *C. borealis*.

| Gene Annotation Metrics (A) |  | Gene Product Categories (B) |  |
| --- | --- | --- | --- |
| Metric | Value | Category | Count |
| Total Genes | 17,055 | Uncharacterized | 5,893 |
| Unique Products | 8,496 | Receptors / Trans-<br>porters | 826 |
| Median Exons/Gene | 6 | Enzymes | 704 |
| Max Exons/Gene | 210 | Zinc Finger Proteins | 386 |
| Mean Exon Length (bp) | 2,164 | Histone Family | 323 |
| Mean Gene Size (bp) | 18,390 | Cuticle / Structural | 198 |
| Max Gene Size (bp) | 1,317,456 | Cell Adhesion | 92 |
| Unique Chromosomes | 52 | Other | 8,633 |

This broad range of gene sizes reflects substantial intronic expansion in many loci. Such long introns are particularly relevant in nervous systems, where regulatory complexity and alternative splicing are central to neuronal identity and function Grabowski (2011); Chen and Maniatis (2013).

### Functional Composition of the Gene Repertoire

Functional annotation reveals a diverse gene complement underlying the biology of *C. borealis*. Uncharacterized genes and long non-coding RNAs constitute the largest category, reflecting lineage-specific genes and limited representation of crustaceans in existing protein databases. Among annotated genes, receptors and transporters, enzymes, transcription factors, structural proteins, and cell adhesion molecules are well represented (Table 2 Gene Product Categories(B)). This diversity provides the molecular substrate for the robust sensory, motor, and modulatory capabilities of the crab nervous system.

### Genome-Wide Identification of Neuronal Gene Families

Genome anchoring of previously identified neuronal transcripts resulted in the confident mapping of 68 neuronal genes spanning 15 neural-related gene families (Table 3). These families include ligand-gated receptors, voltage-gated ion channels, and neuromodulatory receptors critical for synaptic transmission and circuit dynamics.

**Table 3.** Neural-related gene families annotated in *C. borealis*.

| Gene Family | Genes (Mapped) | Avg. Exons | Avg. Exon Length (bp) | Avg. Gene Size (bp) |
| --- | --- | --- | --- | --- |
| Innexin subtypes | 6 (6) | 5.0 | 1,884.8 | 79,652.3 |
| Acetylcholine receptors | 14 (12) | 10.8 | 2,685.7 | 87,033.5 |
| Glutamate-gated Cl<sup>-</sup> channel | 2 (1) | 10.0 | 1,467.0 | 17,308.0 |
| Glutamate receptors | 18 (13) | 14.8 | 3,522.1 | 89,215.4 |
| GABA receptors | 8 (6) | 9.7 | 1,935.2 | 32,606.3 |
| Histamine receptors | 3 (3) | 9.7 | 1,518.3 | 21,797.3 |
| Serotonin receptors | 4 (1) | 5.0 | 2,208.0 | 17,848.0 |
| Dopamine receptors | 3 (2) | 3.5 | 3,324.5 | 80,038.0 |
| Octopamine / Tyramine receptors | 6 (5) | 3.4 | 1,877.8 | 40,704.0 |
| Transient receptor potential channels | 6 (6) | 23.3 | 4,313.5 | 34,759.0 |
| Hyperpolarization-activated channels | 5 (5) | 14.0 | 3,946.6 | 66,786.6 |
| Na<sup>+</sup> channels | 2 (2) | 30.5 | 5,713.0 | 43,820.0 |
| Ca<sup>2+</sup> channels | 3 (2) | 39.5 | 8,590.5 | 113,288.0 |
| Other K<sup>+</sup> channels | 7 (7) | 10.7 | 2,946.4 | 166,529.4 |
| Voltage-dependent K<sup>+</sup> channels | 10 (10) | 13.3 | 4,473.0 | 135,981.5 |

Ionotropic glutamate receptors represent the largest family (18 genes), followed by acetylcholine receptors (14 genes) and voltage-dependent potassium channels (10 genes). Sodium and calcium channels are fewer in number, but exhibit exceptional structural complexity, with exon counts exceeding 30–40 per gene.

The ionotropic glutamate receptor family represents the largest neuronal gene class identified in the *Cancer borealis* genome, comprising 18 genes, followed by acetylcholine receptors (14 genes) and voltage-dependent potassium channels (10 genes) (Table 3). Although fewer in number, calcium and sodium channels exhibit particularly complex exon architectures, with some loci exceeding 40 exons. Together, these gene families illustrate the extensive molecular diversity underlying neuronal signaling in *C. borealis*, reinforcing its value as a model system for studying neural circuits and synaptic physiology.

### Genome-Anchored Architecture of Neuronal Gene Families

The following subsections describe the genomic organization of each major neuronal gene family using genome-anchored annotation derived from Table 4.

**Table 4.** Summary of Transcriptome-Annotated Neuronal Genes in *Cancer borealis* (de novo assembly and mapping with STAR aligner)

| Accession | Symbol | Exons | Exon(bp) | Ch Contig No | Chr No. | Span | Start | End | DNASTAR | STAR | Trinity |
| --- | --- | --- | --- | --- | --- | --- | --- | --- | --- | --- | --- |
| **Innexin Subtypes** |  |  |  |  |  |  |  |  |  |  |  |
| JQ994479.1 | INX1 | 7 | 1234 | CM084938.1 | chr 26 | 6561 | 10382089 | 10388649 | 62610 | 132542 | 226645 |
| JQ994480.1 | INX2 | 5 | 1451 | CM084938.1 | chr 26 | 3709 | 10306096 | 10309804 | 96143 | 261089 | 303873 |
| JQ994481.1 | INX3 | 4 | 1264 | CM084918.1 | chr 6 | 32487 | 7791138 | 7823624 | 19865 | 47276 | 130662 |
| KJ642222.1 | INX4 | 4 | 2833 | CM084922.1 | chr 10 | 423346 | 12277169 | 12700514 | 2 | 16 | 1139 |
| KJ817410.1 | INX5 | 5 | 1074 | CM084938.1 | chr 26 | 2594 | 10313957 | 10316550 | 7338 | 19536 | 16850 |
| KJ817411.1 | INX6 | 5 | 3209 | CM084914.1 | chr 2 | 9217 | 532 | 2875 | 532 | 2875 | 3905 |
| **Acetylcholine Receptors** |  |  |  |  |  |  |  |  |  |  |  |
| KX021822 | mAChR-A | NO_HIT | - | - | - | - | - | - | 13 | - | 1983 |
| KX021821 | mAChR-B | NO_HIT | - | - | - | - | - | - | 23 | - | 1871 |
| KX021828 | nAChR-alpha1 | 6 | 4077 | CM084947.1 | chr 35 | 106827 | 4977459 | 5084285 | 6 | 32 | 4924 |
| KX021827 | nAChR-alpha2 | 9 | 1960 | CM084947.1 | chr 35 | 131634 | 5153051 | 5284684 | 20 | 32 | 397 |
| KX021829 | nAChR-alpha3 | 10 | 1281 | CM084941.1 | chr 29 | 11568 | 6789613 | 6801180 | 26 | 46 | 6359 |
| KX021830 | nAChR-alpha4 | 18 | 3117 | CM084923.1 | chr 11 | 67518 | 9691873 | 9759390 | 3929 | 30 | 6359 |
| KX021824 | nAChR-alpha5 | 9 | 1452 | CM084934.1 | chr 22 | 16332 | 12574094 | 12590425 | 163 | 85 | 432 |
| KX021825 | nAChR-alpha7 | 10 | 1569 | CM084928.1 | chr 16 | 335225 | 6101061 | 6436285 | 18 | 38 | 1206 |
| KX021831 | nAChR-alpha8 | 8 | 5471 | CM084947.1 | chr 35 | 38107 | 5290816 | 5328922 | 93 | 479 | 4924 |
| KX021826 | nAChR-alpha16 | 12 | 1851 | CM084913.1 | chr 1 | 134335 | 37419322 | 37553656 | 26 | 38 | 430 |
| KX021823 | nAChR-beta1 | 12 | 1581 | CM084928.1 | chr 16 | 85663 | 6696638 | 6782300 | 76 | 134 | 2920 |
| **Glutamate-Gated Chloride Channel** |  |  |  |  |  |  |  |  |  |  |  |
| KX059698 | Glu-Cl | 10 | 1467 | CM084914.1 | chr 2 | 17308 | 23586296 | 23603603 | 98 | 77 | 3812 |
| **Glutamate Receptors** |  |  |  |  |  |  |  |  |  |  |  |
| KU986879 | mGluR1 | 16 | 2874 | CM084929.1 | chr 17 | 8678 | 12826979 | 12835656 | 888 | 2272 | 2689 |
| KU986880 | mGluR2 | 18 | 5703 | CM084943.1 | chr 31 | 105013 | 4963936 | 5068948 | 0 | 32 | 831 |
| KU986881 | mGluR3 | 11 | 2682 | CM084945.1 | chr 33 | 55742 | 7528683 | 7584424 | 589 | 1225 | 566 |
| KU986882 | mGluR4 | 18 | 5703 | CM084943.1 | chr 31 | 105013 | 4963936 | 5068948 | 6 | 32 | 831 |
| KU986883 | mGluR5 | 23 | 5388 | CM084916.1 | chr 4 | 23723 | 27659724 | 27683446 | 9973 | 2169 | 4571 |
| KU986884 | mGluR7 | NO_HIT | - | - | - | - | - | - | 151 | - | 1697 |
| **GABA Receptors** |  |  |  |  |  |  |  |  |  |  |  |
| KU986868 | mGABAr-1 | 19 | 2667 | CM084945.1 | chr 33 | 13696 | 6454633 | 6468328 | 1798 | 783 | 3160 |
| KU986869 | mGABAr-2 | NO_HIT | - | - | - | - | - | - | 141 | - | 1550 |
| KU986871 | LCCH3-like | 8 | 1446 | CM084924.1 | chr 12 | 78386 | 14933037 | 15011422 | 425 | 825 | 4059 |
| KU986872 | RDL-like | 11 | 2240 | CM084945.1 | chr 33 | 62925 | 1333795 | 1396719 | 17 | 75 | 3193 |
| KU986873 | GRD-like | 12 | 1770 | CM084927.1 | chr 15 | 13785 | 12908105 | 12921889 | 107 | 169 | 598 |
| **Histamine Receptors** |  |  |  |  |  |  |  |  |  |  |  |
| KU716100 | HisR1 | 10 | 1491 | CM084945.1 | chr 33 | 29661 | 7636302 | 7665962 | 62 | 123 | 1086 |
| KU716101 | HisR2 | 10 | 1491 | CM084945.1 | chr 33 | 29661 | 7636302 | 7665962 | 35 | 123 | 121 |
| KU716102 | HisR3 | 9 | 1573 | CM084955.1 | chr 43 | 6070 | 2943271 | 2949340 | 77 | 190 | 615 |
| **Serotonin Receptors** |  |  |  |  |  |  |  |  |  |  |  |
| KU710381 | HTR1A | NO_HIT | - | - | - | - | - | - | 2493 | - | 1419 |
| KU710382 | HTR1B | NO_HIT | - | - | - | - | - | - | 128 | - | 922 |
| KU710380 | HTR2B |  |  |  |  |  |  |  | 13 |  | 2735 |
| KU710379 | HTR7 | 5 | 2208 | CM084917.1 | chr 5 | 7848 | 5064603 | 5082450 | 52 | 387 | 18104 |
| **Dopamine Receptors** |  |  |  |  |  |  |  |  |  |  |  |
| KU710377 | D1αR | 4 | 5398 | CM084933.1 | chr 21 | 148573 | 3798143 | 3946715 | 13 | 80 | 945 |
| KU710376 | D1βR | NO_HIT | - | - | - | - | - | - | 27 | - | 1565 |
| KU710378 | D2αR | 8 | 1800 | CM084914.1 | chr 2 | 505133 | 19764367 | 20269499 | 238 | 460 | 1272 |
| **Octopamine/Tyramine Receptors** |  |  |  |  |  |  |  |  |  |  |  |
| KU710373 | Tyr-R | 1 | 1401 | CM084928.1 | chr 16 | 1401 | 3026213 | 3027613 | 97 | 194 | 933 |
| KU710375 | OctαR | 1 | 495 | CM084921.1 | chr 9 | 495 | 8982448 | 8982942 | 0 | 0 | 102 |
| KU710372 | OctβR1 | 11 | 3858 | CM084921.1 | chr 9 | 116949 | 3499482 | 3616430 | 126 | 282 | 1018 |
| KU710374 | OctβR2 | NO_HIT | - | - | - | - | - | - | 98 | - | 1577 |

| Accession | Symbol | Exons | Exon(bp) | Ch Contig No | Chr No. | Span | Start | End | DNASTAR | STAR | Trinity |
| --- | --- | --- | --- | --- | --- | --- | --- | --- | --- | --- | --- |
| KU710370 | OctβR3 | 2 | 1823 | CM084942.1 | chr 30 | 18811 | 1695862 | 1714672 | 6 | 17 | 1538 |
| KU710371 | OctβR4 | 2 | 1823 | CM084930.1 | chr 18 | 36186 | 13996892 | 14033077 | 8386 | 19720 | 3060 |
| **Transient Receptor Potential (TRP)** |  |  |  |  |  |  |  |  |  |  |  |
| KX037435 | TRP-A1 | 24 | 3648 | CM084947.1 | chr 35 | 25011 | 1549076 | 1574086 | 7189 | 82 | 96 |
| KX037434 | TRP-A-like | 26 | 3675 | CM084913.1 | chr 1 | 31921 | 27185578 | 27217498 | 11 | 2978 | 3629 |
| KX037436 | TRP-M1 | 23 | 5205 | CM084919.1 | chr 7 | 17166 | 24706978 | 24724143 | 292 | 1063 | 1823 |
| KX037433 | TRP-M3 | 23 | 5205 | CM084919.1 | chr 7 | 17166 | 24706978 | 24724143 | 108 | 1063 | 1823 |
| KX037437 | TRP-M-like | 23 | 5205 | CM084919.1 | chr 7 | 17166 | 24706978 | 24724143 | 87 | 1063 | 1823 |
| KX037438 | TRP-V5 | 21 | 2943 | CM084936.1 | chr 24 | 100124 | 3804073 | 3904196 | 0 | 9 | 12 |
| **Hyperpolarization-Activated and cyclic Nucleotide-Gated Channels** |  |  |  |  |  |  |  |  |  |  |  |
| DQ103257 | HCN/IH | 10 | 2007 | CM084925.1 | chr 13 | 252083 | 14939827 | 15191909 | 182 | 332 | 7957 |
| KU716097 | CNG-Alpha1 | 5 | 2830 | CM084961.1 | chr 49 | 9663 | 754714 | 764376 | 213 | 212 | 764 |
| KU716098 | CNG-Alpha2 | 13 | 4405 | CM084947.1 | chr 35 | 9473 | 5794928 | 5804400 | 149 | 414 | 1455 |
| KU716099 | CNG-Alpha3 | 21 | 4971 | CM084961.1 | chr 49 | 49905 | 697166 | 747070 | 51 | 229 | 602 |
| KU716096 | CNG-Beta1 | 21 | 5520 | CM084930.1 | chr 18 | 12809 | 4000942 | 4013750 | 1065 | 4938 | 10183 |
| **Voltage-dependent K<sup>+</sup> Channels** |  |  |  |  |  |  |  |  |  |  |  |
| FJ263946 | shaker | 7 | 1686 | CM084923.1 | chr 11 | 114765 | 20051572 | 20166336 | 68 | 186 | 3618 |
| DQ103255 | shab | 8 | 3387 | CM084947.1 | chr 35 | 54945 | 6969881 | 7024825 | 58 | 152 | 1944 |
| KU681456 | shaw1 | 7 | 7019 | CM084930.1 | chr 18 | 72851 | 845367 | 918217 | 10 | 131 | 1863 |
| KU681455 | shaw2 | 13 | 2232 | CM084926.1 | chr 14 | 28691 | 9930251 | 9958941 | 82 | 133 | 1108 |
| DQ103254 | shal | 6 | 2025 | CM084915.1 | chr 3 | 82099 | 25420416 | 25502514 | 28 | 101 | 2432 |
| KU681453 | KCNQ1 | 17 | 4565 | CM084938.1 | chr 26 | 141525 | 4018493 | 4160017 | 150 | 2395 | 26828 |
| KU681452 | KCNQ2 | 15 | 7239 | CM084923.1 | chr 11 | 645221 | 2978842 | 3624062 | 33 | 110 | 2002 |
| KU681458 | KCNH1/EAG | 22 | 6355 | CM084947.1 | chr 35 | 140778 | 4529233 | 4670010 | 41 | 609 | 4962 |
| KU681459 | KCNH2 | 12 | 3234 | CM084917.1 | chr 5 | 18254 | 9559513 | 9577766 | 5 | 14 | 2452 |
| KU681460 | KCNH3 | 26 | 6988 | CM084940.1 | chr 28 | 60686 | 2181101 | 2241786 | 141 | 752 | 75689 |
| **Other K<sup>+</sup> channels** |  |  |  |  |  |  |  |  |  |  |  |
| DQ103256 | BKCa | 25 | 3138 | CM084942.1 | chr 30 | 91529 | 8046956 | 8138484 | 922 | 2036 | 26485 |
| KU710383 | SKCa | 11 | 7271 | CM084922.1 | chr 10 | 512663 | 11389572 | 11902234 | 2022 | 1240 | 3600 |
| KU681454 | KCNT1 | 31 | 4830 | CM084923.1 | chr 11 | 445839 | 18074517 | 18520355 | 76 | 200 | 3039 |
| KU681451 | IRK | 10 | 1920 | CM084938.1 | chr 26 | 64250 | 908181 | 972430 | 4 | 12 | 797 |
| KU681438 | KCNK1 | 8 | 5297 | CM084920.1 | chr 8 | 467650 | 11238324 | 11705973 | 34 | 301 | 1680 |
| KU681437 | KCNK2 | 7 | 1116 | CM084946.1 | chr 34 | 15139 | 3346401 | 3361539 | 50 | 66 | 268 |
| **Ca<sup>2+</sup> Channels** |  |  |  |  |  |  |  |  |  |  |  |
| KU702651 | CaV1 | 34 | 4047 | CM084938.1 | chr 26 | 81676 | 9533592 | 9615267 | 2923 | 4833 | 19379 |
| JN809808 | CaV2 | 41 | 8697 | CM084946.1 | chr 34 | 116471 | 3092483 | 3208953 | 349 | 2818 | 5426 |
| JN809810 | CaV3 | 38 | 8484 | CM084915.1 | chr 3 | 110105 | 32763209 | 32873313 | 47 | 360 | 10813 |
| **Na<sup>+</sup> Channels** |  |  |  |  |  |  |  |  |  |  |  |
| EF089568 | NaV | 30 | 6206 | CM084932.1 | chr 20 | 71275 | 2930994 | 3002268 | 1114 | 2263 | 23557 |
| KU681457 | NALCN | 31 | 5220 | CM084915.1 | chr 3 | 16365 | 33088509 | 33104873 | 832 | 1607 | 4914 |
| **Kainate-Like Receptors** |  |  |  |  |  |  |  |  |  |  |  |
| KX016772 | Kainate-1A | 6 | 1183 | CM084945.1 | chr 33 | 311814 | 1403899 | 1715712 | 45 | 22 | 1113 |
| KX016773 | Kainate-1B | 19 | 2589 | CM084936.1 | chr 24 | 44150 | 2005648 | 2049797 | 7 | 51 | 914 |
| KX016774 | Kainate-2A | 20 | 7083 | CM084944.1 | chr 32 | 73957 | 5894255 | 5968211 | 128 | 376 | 2304 |
| KX016775 | Kainate-2B | 22 | 3668 | CM084919.1 | chr 7 | 172680 | 3997353 | 4170032 | 466 | 2054 | 53924 |
| KX016776 | Kainate-2C | 19 | 2589 | CM084936.1 | chr 24 | 44150 | 2005648 | 2049797 | 113 | 51 | 944 |
| **NMDA-like Receptors** |  |  |  |  |  |  |  |  |  |  |  |
| KX016782 | NMDA-1A | 19 | 4823 | CM084929.1 | chr 17 | 37554 | 11056651 | 11094204 | 329 | 897 | 19906 |
| KX016783 | NMDA-1B | NO_HIT | - | - | - | - | - | - | 37 | - | 3 |
| KX016785 | NMDA-2A | 16 | 2602 | CM084940.1 | chr 28 | 26284 | 91622 | 117905 | 2057 | 451 | 5568 |
| KX016786 | NMDA-2B | NO_HIT | - | - | - | - | - | - | 192 | - | 5047 |
| KX016784 | NMDA-2-like | 20 | 2877 | CM084928.1 | chr 16 | 118252 | 13452233 | 13570484 | 81 | 160 | 1668 |

**Table 5.** Mapping statistics of the *de novo* assembled transcriptome against the *Cancer borealis* reference genome.

| Alignment Metric | Count | Percentage |
| --- | --- | --- |
| Total Assembled Transcripts | 330,516 | 100.00% |
| Primary Mapped Transcripts | 280,084 | 84.74% |
| Secondary Alignments (Isoforms) | 125,680 | — |
| Supplementary Alignments | 22,516 | — |
| Unmapped Transcripts | 50,432 | 15.26% |

Note: The overall mapped alignments exceed the total transcript count because a single transcript can produce secondary alignments if it maps to multiple highly similar regions in the genome.

#### Innexin Subtypes

Six innexin genes were identified and genome-anchored in *Cancer borealis*, encoding the gap junction proteins responsible for electrical coupling in invertebrate nervous systems. This gene count is fully consistent with previous molecular and transcriptomic studies of the stomatogastric nervous system, which reported six innexin subtypes expressed in *C. borealis* neurons Shruti *et al*. (2014); Northcutt *et al*. (2016). The concordance between genome-based annotation and prior expression-based analyses provides independent validation of the completeness and accuracy of the current annotation. The six innexins exhibit modest exon numbers (4–7 exons) but differ substantially in their genomic spans, reflecting variation in intronic content. Three innexin genes (INX1, INX2, and INX5) are localized to chromosome 26, suggesting localized clustering that may reflect tandem duplication or shared regulatory architecture. The remaining innexins are distributed across chromosomes 2, 6, and 10 (Table 4). Such chromosomal organization is consistent with the functional role of innexins in forming electrically coupled networks, where coordinated regulation of gap junction expression is critical for synchronous neuronal activity.

Previous physiological and molecular studies demonstrated that differential innexin expression contributes to neuron-specific electrical coupling patterns within the stomatogastric ganglion Shruti *et al*. (2014). The genome-anchored definition of innexin loci presented here extends these findings by revealing the underlying gene structures and chromosomal context, providing a framework for future studies investigating regulatory elements, alternative splicing, and evolutionary conservation of electrical synapse components in crustacean neural circuits.

#### Acetylcholine Receptors

Fourteen acetylcholine receptor genes were identified in the *Cancer borealis* genome, including multiple nicotinic acetylcholine receptor (nAChR) subunits and two muscarinic acetylcholine receptors. Genome anchoring revealed substantial heterogeneity in gene structure, chromosomal location, and duplication status across this family (Table 4). Multiple nicotinic receptor subunits that were previously represented by single transcriptome-derived sequences map to distinct chromosomal loci, providing clear evidence of gene duplication. For example, nAChR-*β*1 and nAChR-*α*4 each map to separate chromosomes, consistent with duplication followed by divergence. Such duplication events are common in ligand-gated ion channel families and are thought to facilitate functional diversification through altered expression patterns, subunit composition, and pharmacological sensitivity Ohno (1970); Lynch and Conery (2000).

In addition to duplication across chromosomes, localized clustering of acetylcholine receptor genes is observed. Several nAChR subunits are co-localized on chromosome 35, forming a small genomic cluster analogous to the innexin cluster identified on chromosome 26. Gene clustering within neurotransmitter receptor families has been reported in both invertebrates and vertebrates and may reflect tandem duplication events or selection for shared regulatory control Ortells and Lunt (1997); Le Novère and Changeux (2002). Such organization can enable coordinated transcriptional regulation of receptor subunits that function together within synaptic complexes. By contrast, both muscarinic acetylcholine receptors (mAChR-A and mAChR-B) remain unassigned to chromosome-length scaffolds. These loci likely reside on unplaced contigs or represent structurally complex regions that are not yet fully resolved in the current assembly. The presence of muscarinic receptor transcripts in prior transcriptomic studies indicates that these genes are expressed but underscores the continuing need for assembly refinement to resolve all components of cholinergic signaling in *C. borealis*.

#### Glutamate-Gated Chloride Channel (Glu-Cl)

A single glutamate-gated chloride channel gene (Glu-Cl; KX059698) was identified and mapped to chromosome 2 (CM084914.1). This gene contains 10 exons with a total exon length of 1,467 bp and spans approximately 17.3 kb of genomic sequence. Although only a single copy was detected, the gene exhibits a genomic span substantially larger than its coding sequence, indicating extensive intronic regions. Glu-Cl channels play a central role in inhibitory neurotransmission in many invertebrate nervous systems and are subject to complex regulatory control Dent (2000). The presence of long introns may facilitate regulatory flexibility, including alternative splicing or developmental and cell-type-specific expression. Unlike nicotinic acetylcholine or glutamate receptors, Glu-Cl appears not to have undergone recent duplication in *C. borealis*, suggesting stronger selective constraints on gene dosage or channel composition in this inhibitory pathway.

#### Glutamate Receptors

Eighteen glutamate receptor genes were identified, representing ionotropic NMDA-like and kainate-like receptors as well as metabotropic glutamate receptors (mGluRs). This family displays extensive structural and evolutionary diversity, with exon counts ranging from as few as six to more than twenty exons per gene. Several glutamate receptors show clear evidence of gene duplication, with paralogous loci mapping to distinct chromosomes (Table 4). For example, Kainate-2A and NMDA-2-like receptors are each represented by multiple genomic loci, consistent with duplication events followed by divergence. Duplication within glutamate receptor families has been widely documented and is thought to support the expansion of synaptic signaling modalities, kinetic diversity, and plasticity across neural circuits Traynelis *et al*. (2010). Such paralogous receptors may differ in expression patterns, intracellular coupling, or pharmacological properties, enabling fine-tuning of excitatory transmission. In addition to dispersed duplication, partial clustering of glutamate receptor genes is observed on specific chromosomes, suggesting a history of tandem duplication and local retention. Similar genomic organization has been reported in other metazoans and may facilitate coordinated regulation of functionally related receptor subtypes Greer and Greenberg (2004).

#### GABA Receptors

Eight GABA receptor genes were identified and examined in the *Cancer borealis* genome, including both ionotropic and metabotropic subtypes that mediate inhibitory neurotransmission. This gene count is consistent with previous transcriptomic analyses of the stomatogastric nervous system, which identified a comparable complement of GABA receptor transcripts Northcutt *et al*. (2016). Genome anchoring confirms the majority of these loci and provides insight into their structural organization and evolutionary history. A particularly notable feature of this gene family is the GRD-like GABA receptor, which maps to multiple distinct chromosomal loci, providing strong evidence for gene duplication. Such duplication events within inhibitory receptor families have been reported in other invertebrate lineages and are thought to contribute to functional diversification through alterations in expression pattern, channel kinetics, or synaptic localization Lynch and Conery (2000); Innan and Kondrashov (2010). In neuronal circuits, the presence of paralogous GABA receptor subunits may support flexible inhibitory control across different neuron types or developmental stages.

Other GABA receptor subtypes, including RDL-like and LCCH3-like receptors, exhibit moderate exon counts and comparatively compact coding regions, suggesting more conserved structural organization. In contrast, the metabotropic GABA receptor mGABAr-2 remains unassigned to a chromosome-length scaffold. Together, these findings highlight both conserved and evolving components of inhibitory signaling architecture in *C. borealis*.

#### Histamine Receptors

Three histamine receptor genes were identified and mapped to chromosomes 33 and 43. These genes encode ionotropic chloride channels that mediate inhibitory synaptic transmission in arthropods and are functionally distinct from vertebrate histamine receptors Hardie (1987). All three histamine receptors exhibit consistent exon architectures, containing nine to ten exons, and relatively modest genomic spans. The conserved gene structure observed among histamine receptors suggests strong functional constraint, consistent with their essential role in sensory processing and neural inhibition. Similar architectural conservation has been reported in other insects and crustaceans, indicating that histaminergic signaling pathways are evolutionarily stable across arthropods Gisselmann *et al*. (2002).

#### Serotonin Receptors

Four serotonin receptor transcripts were examined; however, only one receptor (HTR7) was successfully mapped to chromosome 5. HTR7 contains five exons and displays a compact coding structure relative to its genomic span. The remaining serotonin receptors (HTR1A and HTR1B) were not assigned to chromosome-length scaffolds. This partial genomic resolution mirrors patterns observed in earlier transcriptome-based studies and likely reflects localization on unplaced contigs or increased sequence divergence relative to available reference models Northcutt *et al*. (2016). Serotonin receptors are key neuromodulators implicated in circuit plasticity and state-dependent control of motor output in crustaceans. Resolving the genomic context of the remaining serotonin receptor subtypes will be important for understanding regulatory mechanisms underlying serotonergic modulation in the stomatogastric nervous system.

#### Dopamine Receptors

Three dopamine receptor genes were identified in the *C. borealis* genome. Two receptors, D1*α*R and D2*α*R, were successfully mapped to chromosomes 21 and 2, respectively. The third receptor, D1*β*R, remains unassigned to a chromosome-length scaffold. The mapped dopamine receptors display relatively compact exon architectures but occupy large genomic spans, indicating substantial intronic content. Such organization suggests that regulatory elements within introns may play an important role in controlling receptor expression. Dopamine is a well-characterized neuro-modulator in crustacean neural circuits, where it exerts cell-specific and state-dependent effects on neuronal excitability Harris-Warrick (2010). The presence of multiple dopamine receptor subtypes, coupled with evidence of regulatory complexity at the genomic level, supports a model in which dopaminergic signaling achieves functional diversity through differential gene regulation rather than extensive expansion of receptor number.

#### Octopamine and Tyramine Receptors

Six octopamine and tyramine receptor genes were identified and genome-anchored in *Cancer borealis*, representing key components of biogenic amine signaling pathways in crustacean nervous systems. These receptors are distributed across multiple chromosomes, indicating an absence of strong genome-wide clustering, although limited local proximity is observed for some subtypes (Table 4). Most octopamine and tyramine receptors exhibit low exon counts, often comprising one to a few exons, yet their genomic spans vary substantially, suggesting differences in intronic content and regulatory architecture. Such organization is consistent with G-protein-coupled receptor (GPCR) families, where regulatory elements rather than extensive exon multiplication often underlie functional specialization Hauser *et al*. (2006). Gene duplication within biogenic amine receptor families has been reported across arthropods and is thought to facilitate the diversification of modulatory effects on neural circuits, including changes in intracellular signaling pathways and neuron-specific expression patterns Roeder (2005).

#### Transient Receptor Potential (TRP) Channels

Six transient receptor potential (TRP) channel genes were mapped in the *C. borealis* genome, representing some of the most structurally complex neuronal genes identified in this study. These genes contain among the highest exon counts observed, ranging from 21 to 26 exons, and are distributed across several chromosomes. Their large genomic spans indicate extensive intronic regions (Table 4). TRP channels are polymodal sensory channels implicated in mechanosensation, thermosensation, and chemosensation, and their complex exon– intron architectures are consistent with roles in fine-tuned sensory integration Clapham (2003). The presence of long introns may provide regulatory flexibility, enabling cell-type-specific expression or alternative splicing.

#### Hyperpolarization-Activated and Cyclic Nucleotide-Gated Channels

Five genes encoding hyperpolarization activated (HCN) and cyclic nucleotide-gated (CNG) channels were identified and mapped to chromosomes 13, 18, 35, and 49. These channels contain between 10 and 21 exons and occupy moderate genomic spans relative to their coding lengths. Such structural complexity is consistent with their integrative roles in regulating membrane excitability, rhythmic activity, and responsiveness to neuromodulators. HCN and CNG channels serve as key determinants of neuronal pacing and resonance, particularly in central pattern-generating circuits Biel *et al*. (2009). The observed exon diversity may support regulatory control through alternative splicing or promoter usage, allowing neurons to fine-tune intrinsic electrical properties in response to developmental or modulatory cues.

#### Na^+^ and Ca^2+^ Channels

Voltage-gated sodium and calcium channels exhibit the greatest exon complexity among all neuronal gene families examined. The voltage-gated sodium channel (NaV) and the sodium leak channel (NALCN) contain 30 and 31 exons, respectively, while the calcium channels CaV2 and CaV3 contain 41 and 38 exons (Table 4). Despite their limited number, these genes span unusually large genomic regions. Such extensive exon–intron architectures are characteristic of channels that perform essential roles in action potential initiation, propagation, and calcium-dependent intracellular signaling Catterall (2011). Large introns may mediate fine spatiotemporal regulation of channel expression critical for neuronal physiology Santin and Schulz (2019).

#### Potassium Channels

Potassium channels in *C. borealis* span both voltage-dependent and non–voltage-dependent subfamilies and display wide variation in genomic organization. Several channels, including KCNK1, KCNK2, and KCNT1, exhibit disproportionately large genomic spans relative to their total exon lengths, indicating extensive intronic regions. In contrast, members of the Shaker family retain compact exon counts while occupying large genomic regions. This pattern reflects a decoupling of protein sequence conservation from genomic organization. Shaker-family potassium channels are highly conserved at the amino acid level across decapod crustaceans, yet their intronic architectures vary substantially. Such variation suggests rapid evolution of regulatory elements within non-coding regions, enabling species- or circuit-specific tuning of neuronal excitability without altering fundamental channel properties Marder and Taylor (2011); Goaillard and Marder (2009). These features underscore the importance of regulatory evolution in maintaining functional robustness alongside molecular diversity.

### Structural Diversity of Neuronal Genes

Neuronal genes in *C. borealis* display striking heterogeneity in genomic architecture. Exon number, exon size, and genomic span vary extensively both within and among gene families. Transient receptor potential (TRP) channels, calcium channels, and sodium channels exhibit particularly complex exon–intron structures, whereas several neuromodulatory receptors possess relatively compact coding sequences embedded within large genomic spans. This structural diversity likely reflects differing regulatory demands and evolutionary histories across neuronal signaling pathways.

### Chromosomal Distribution and Localized Clustering

Neuronal genes are broadly distributed across the 51 chromosome-length scaffolds (Figure 2), indicating the absence of extreme genome-wide clustering. Nevertheless, localized clustering is evident within certain families, such as innexins on chromosome 26 and multiple acetylcholine receptor subunits on chromosome 35. Such clustering may result from tandem duplication events or shared regulatory environments, consistent with observations in other metazoan genomes Lynch and Conery (2000).

**Figure 2.**
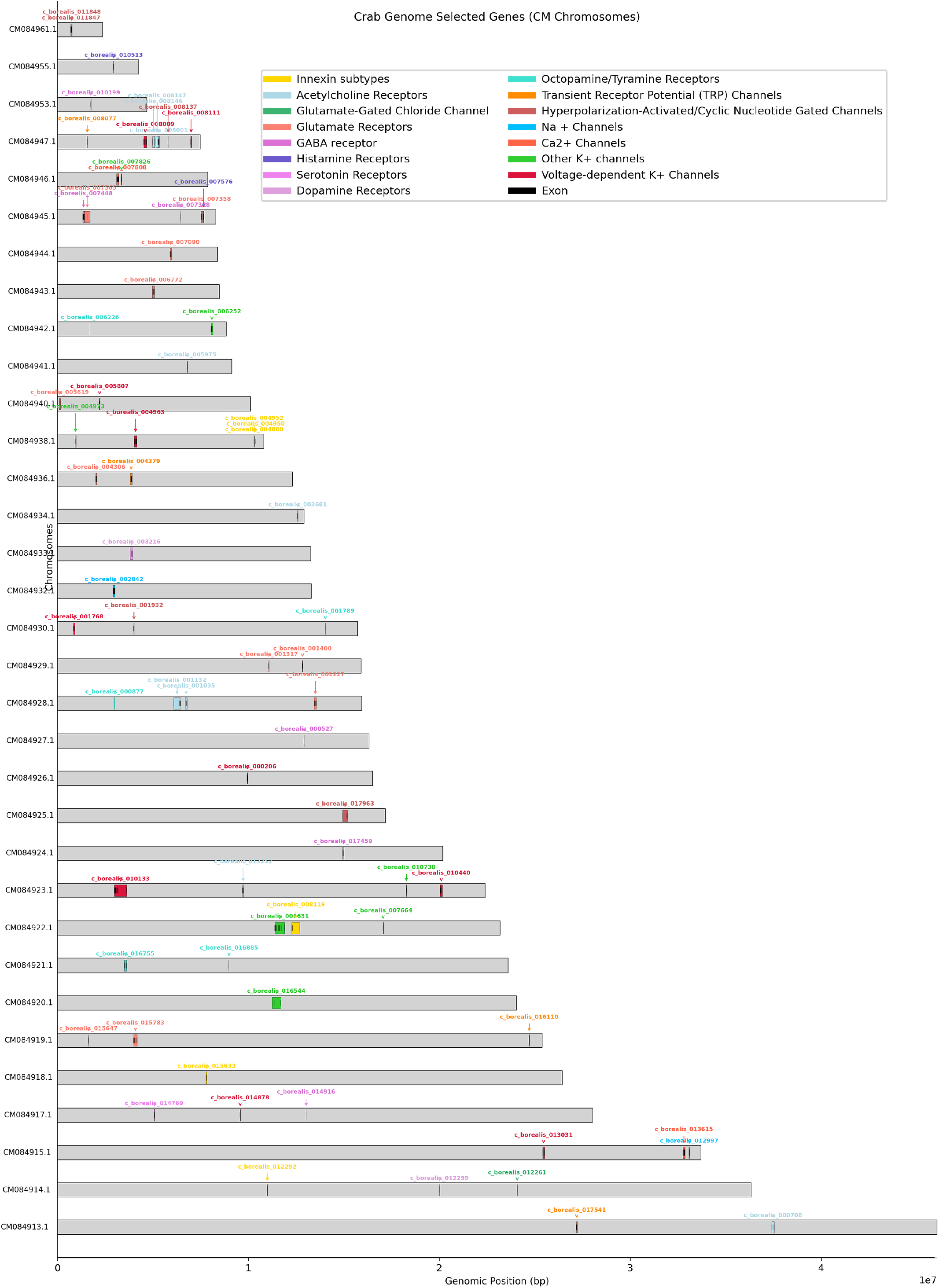
Chromosomal distribution and relative genomic spans of neuronal genes in *Cancer borealis*. Each bar represents a chromosome-length scaffold, and colored segments within bars indicate the chromosomal location and relative genomic span of annotated neuronal genes. Colors correspond to distinct neuronal gene families, as shown in the legend.

### Gene Duplication as a Driver of Neuronal Diversity

A central finding of genome-based annotation is the identification of multiple genomic loci corresponding to transcripts previously interpreted as single genes. This pattern is particularly evident in acetylcholine receptors, glutamate receptors, and GABA receptors. For example, nAChR-*β*1 and nAChR-*α*4 each map to distinct chromosomal loci, while the GRD-like GABA receptor is represented by multiple paralogous loci across different chromosomes (Table 4).

Gene duplication is a fundamental evolutionary mechanism that generates genetic redundancy, allowing one gene copy to maintain essential functions while others diverge through subfunctionalization or neofunctionalization Ohno (1970); Innan and Kondrashov (2010). In nervous systems, duplicated ion channels and receptors can evolve distinct expression patterns, kinetics, or pharmacological properties, enabling fine-tuned modulation of circuit behavior Marder (2012); Goaillard and Marder (2009). Such diversification is especially relevant in crustaceans, where functionally similar neurons often exhibit highly variable ion channel expression profiles while maintaining stable output Marder and Taylor (2011).

The genome-anchored identification of paralogous neuronal genes in *C. borealis* therefore provides a mechanistic framework for understanding how conserved circuit function can be achieved through molecular diversity.

### Intronic Expansion and Regulatory Evolution

Comparison of total exon length with overall gene span highlights the substantial contribution of intronic regions to neuronal gene architecture. Potassium channels such as KCNK1, KCNK2, and members of the Shaker family exhibit compact coding sequences embedded within large genomic regions. While amino acid sequences of these channels are highly conserved across decapods, intron number and size vary considerably.

This decoupling of protein conservation and genomic organization suggests that regulatory evolution is occurring primarily at the level of non-coding sequences Keren *et al*. (2010). Large intronic regions can harbor enhancers, silencers, splicing regulators, and non-coding RNAs that modulate gene expression and isoform diversity Chen and Maniatis (2013). Such regulatory flexibility may enable species-specific tuning of neuronal excitability without altering fundamental channel properties.

### Unassigned Neuronal Genes and Assembly Limitations

Despite the quality of the current assembly, a subset of neuronal genes remains unassigned to chromosome-length scaffolds, including muscarinic acetylcholine receptors, certain glutamate and GABA receptor subtypes, serotonin receptors, and specific TRP channels. These loci likely reside on unplaced contigs or represent structurally complex regions that remain difficult to resolve. Continued assembly refinement and targeted annotation will be required to fully capture the neuronal gene repertoire of *C. borealis*.

### Transcriptome Assembly Validation

Alignment of the Trinity *de novo* transcriptome to the *Cancer borealis* reference genome demonstrated a high degree of concordance, confirming the robustness of the assembly. Of the 330,516 total assembled transcripts, 84.74% (280,084 transcripts) successfully mapped to the reference genome as primary alignments. When accounting for secondary and supplementary alignments— which are highly expected due to the mapping of alternative splice variants (isoforms) generated by the Trinity assembler— the overall mapping rate reached 89.47%. This high primary mapping efficiency (> 80%) is indicative of a high-quality transcriptome assembly that accurately captures the transcribed genome of a complex crustacean. The unmapped fraction (~10–15%) likely represents a combination of highly divergent alleles, fragmented untranslated regions (UTRs) that lack sufficient length for confident genomic placement, or expected biological noise inherent to *de novo* assembly algorithms.

### Transcriptome Mapping Validation and Concordance

We compared transcript quantification across three different approaches utilizing reads from RNA-seq analysis from a novel sample of pooled crab STG tissue. We focused on quantification of transcripts for 81 candidate genes important for neuronal function in this circuit including voltage-dependent ion channels, neurotransmitter receptors and gap junction proteins. We compared three distinct methods: mapping of fastq reads to the reference transcriptome generated by Northcutt *et al*. (2016) using the DNASTAR platform, a reference-guided mapping approach with the C. borealis genome assembly as reference using STAR and featureCounts, and a de novo assembly and quantification workflow using Trinity and RSEM.

While there was general concordance in transcript numbers across methods, there was substantially more agreement between the DNASTAR and STAR alignment counts that compare the transcriptome reference Northcutt *et al*. (2016) and our genome assembly (*R* = 0.873, Pearson Correlation) than either of these methods with the Trinity de novo assembly (*R* = 0.618 and *R* = 0.715, DNASTAR and STAR with Trinity respectively).

### Ion Channel Gene Expression in the Stomatogastric Ganglion

Although admittedly overall abundance of a given transcript does not necessarily correlate to the influence of that gene product on cellular function, assaying relative abundance provides fertile ground for exploration of neuronal function. Because the STG is a key circuit model for understanding cellular excitability, emergent properties of network architecture, and neuromodulation Marder and Bucher (2007), we explored four classes of putative gene products from the pooled STG RNAseq data: voltage-dependent ion channels, neurotransmitter and biogenic amine receptors, gap junction proteins and neuropeptide receptors.

Northcutt *et al*. (2016) identified 26 ion channel genes from their analysis of the *C. borealis* neural transcriptome likely involved in neuronal excitability. The counts analysis we performed identified the following voltage-dependent K+ channels as being of the higher abundance transcripts of this group (Figure3): KCNQ1 (Kv7, slow delayed rectifier), KCNH1/EAG (Kv10/ether-a-go-go, non-inactivating delayed rectifier), KCNH3 (Kv12), KCNK1 (K2P1, two-pore leak), BKKCA (KCa1, BK calcium-activated K+), and SKKCA (KCa2, SK calcium-activated K+). Perhaps most strikingly, despite a long history of careful voltage-clamp characterization of ionic currents in STG neurons, virtually none of the ionic currents encoded by homologs of these genes have been isolated and characterized in STG neurons, with the exception of BKKCA Golowasch and Marder (1992); Ransdell *et al*. (2012).

Analysis of calcium and sodium channel counts were more consistent with what has been previously described in STG neurons. Two of three voltage-dependent calcium channels (CaV1, CaV2 but not CaV3) were detected in relatively high abundance (Figure3), consistent with previous characterization of two major calcium currents in crab cells Golowasch and Marder (1992); Ransdell *et al*. (2013). Additionally, the single voltage-dependent sodium channel (NaV/para) and the sodium leak channel NALCN were detected in relatively high abundance. Lastly, the channels responsible for hyperpolarization activated and cyclic nucleotide gated mixed cation channels (HCN/IH and CNG) were all detected at lower but detectable levels (Figure 3).

**Figure 3.**
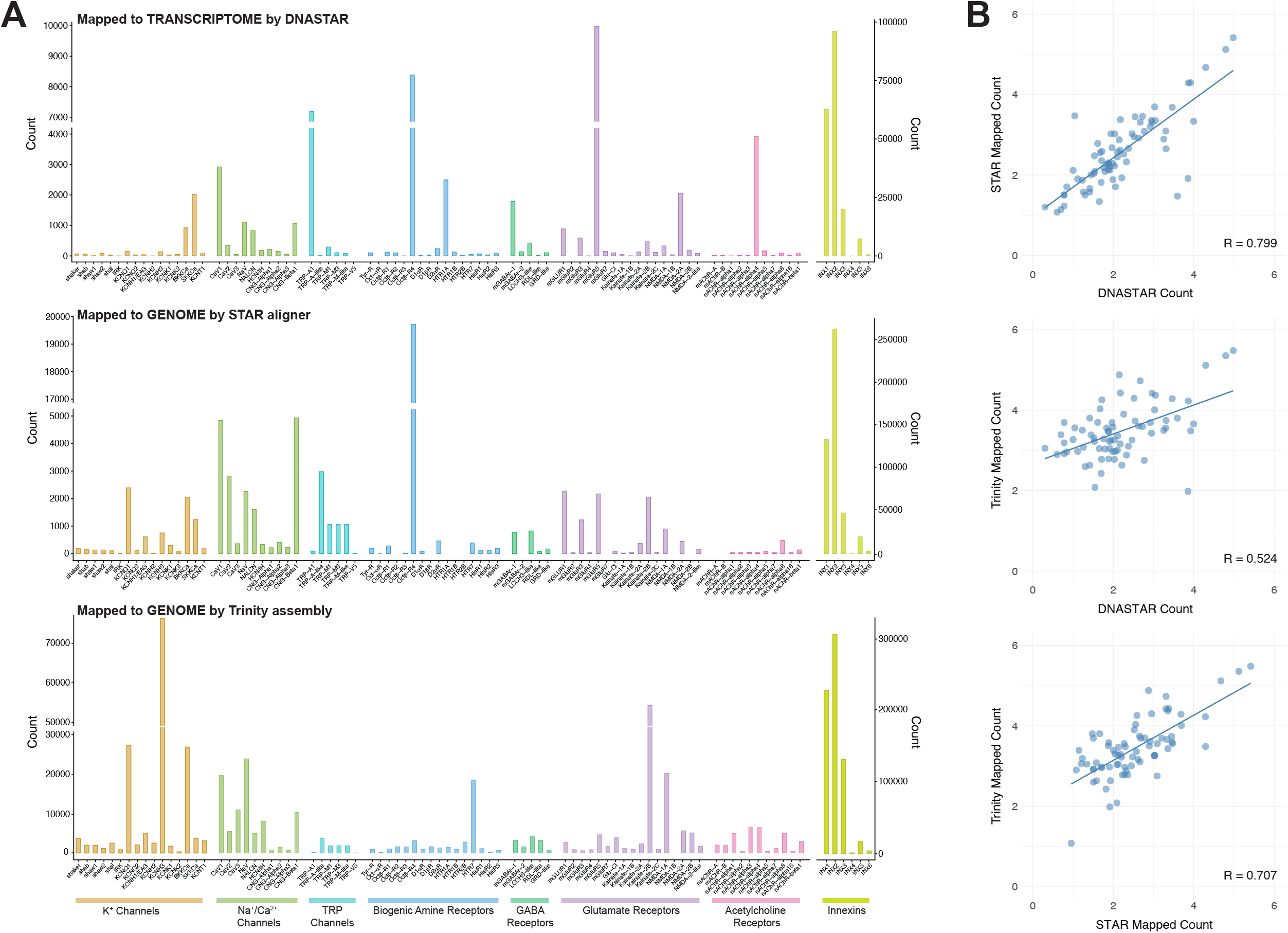
A) Comparison of STG transcript counts for select neuron-related gene families across three estimation techniques: DNAS-tar, STAR aligner and Trinity assembly. Gene families are color coded, and include ion channels, neurotransmitter receptors, and gap junction innexin proteins. B) Comparison of normalized gene expression values (log_10_-transformed counts) among all three estimation approaches. This is a transformation of raw counts, not full CPM normalization. Each point represents a gene. R-value reflects results of Pearson correlation analysis.

Lastly, RNAseq analysis confirms 6 innexin subtypes in the STG Figure 3, with the most predominantly expressed being INX1-3, which has been well studied with focused qPCR analyses in previous work Shruti *et al*. (2014); Northcutt *et al*. (2016).

### Neurotransmitter Receptor Gene Expression in the Stomatogastric Ganglion

A curated set of biogenic amine, glutamate and acetylcholine receptors was generated in the transcriptome analysis of Northcutt *et al*. (2016). Our RNAseq results suggest that octopamine and dopamine each show one predominantly expressed isoform in the STG (Oct*β*4R and D2*α*R respectively; Figure 3). Similarly, one metabotropic (mGABAr1) and one ionotropic (LCCH3-like) GABA receptor show most noticeable expression. However, serotonin receptor expression is less clear: in the DNASTAR mapping to the transcriptome, HTR1A is the clear predominant isoform expressed, while our genome mapping with the STAR aligner did not detect this gene. Rather, HTR7 is the predominant form detected by mapping to the genome. We suggest that finer scale curation of serotonin receptors may be warranted to resolve this more clearly.

The identity of the glutamate and acetylcholine receptors that are responsible for the critical inhibitory synaptic connections throughout the STG have never been formally identified pharmacologically or by other means of analysis. The genome of *C. borealis* contains numerous glutamate receptor isoforms with sequency homology to both NMDA-like and Kainate-like sequences in mammals, as well as a glutamate-gated chloride channel (GluCl). Our RNAseq analysis does not provide any clear candidates for these synaptic receptors, and data from both the DNASTAR and genome aligners are not in concordance. Genome alignment singles out Kainate-2B and NMDA-1A as the strongest candidates by abundance, but other subtypes are clearly detected in the STG transcriptome as well. Similarly, multiple nicotinic isoforms are detected without a clearly predominant form by abundance Figure 3. Given the important role for glutamate and acetylcholine signaling in STG network function Marder and Eisen (1984); Prinz *et al*. (2004), this is a clear area for further analysis and characterization going forward.

### Putative Neuropeptide Receptor Expression in the Stomatogastric Ganglion

Neuropeptide modulation plays a critical role in network function, adaptability and plasticity in the STG. A recent study Fields *et al*. (2026) used a similar genome-mining approach to great effect to characterize many known neuropeptides in *C. borealis*, providing a valuable resource for a genome-enabled framework for discovering endogenous peptide-signaling molecules. However, little is known about the receptors that confer this sensitivity to these broad and numerous number of modulatory peptides Swensen and Marder (2001); Marder (2012). Here we adapt the curation of 46 putative neuropeptide receptors in the *C. borealis* nervous system performed by Dickinson *et al*. (2019) to RNAseq analysis in our pooled STG samples. After mapping reads to the *C. borealis* genome via the STAR aligner, we were able to confirm the presence of 41 of these homologs in the genome and gain an approximation of abundance in the STG (Figure 4). The genomic coordinates for the matching unique genes are listed in Table 6.

**Table 6.** Genomic annotation and expression of neuropeptide precursor genes identified in the *Cancer borealis* genome. Transcript abundance was quantified from adult STG RNA-seq data using the STAR aligner and featureCounts. “None found” indicates that no homologous sequence was identified in the current genome assembly.

| Accession | Neuropeptide | Exons | Exonic Length (bp) | Chromosome | Gene Span (bp) | STAR Count |
| --- | --- | --- | --- | --- | --- | --- |
| GEFB01018628 | Adipokinetic hormone-corazonin-like peptide (ACP) | 6 | 1722 | CM084942.1 | 30031 | 132 |
| GEFB01012018 | Allatostatin A (AST-A) | 9 | 2880 | CM084934.1 | 117681 | 2311 |
| GEFB01014490 | Allatostatin B (AST-B) | 1 | 1197 | CM084925.1 | 1197 | 570 |
| GEFB01019215 | Allatostatin C (AST-C) | – | – | – | – | – |
| None found | Allatotropin | – | – | – | – | – |
| GEFB01007449 | Bursicon (Burs) | 17 | 4200 | CM084940.1 | 33707 | 5463 |
| GEFB01030009 | CCHamide (CCHa) | 3 | 1230 | CM084920.1 | 13961 | 221 |
| GEFB01026704 | Corazonin (CRZ) | – | – | – | – | – |
| GEFB01008615 | Crustacean cardioactive peptide (CCAP) | 9 | 1233 | CM084916.1 | 31925 | 1534 |
| None found | Crustacean hyperglycemic hormone (CHH) | – | – | – | – | – |
| GEFB01018473 | Diuretic hormone 31 (DH31) | 13 | 2076 | CM084939.1 | 132496 | 1639 |
| GEFB01015824 | Diuretic hormone 44 (DH44) | 10 | 1704 | CM084923.1 | 67805 | 483 |
| GEFB01025040 | Ecdysis-triggering hormone (ETH) | 4 | 1611 | CM084929.1 | 21821 | 642 |
| GEFB01000837 | FMRFamide-like peptide (FLP) | 4 | 1281 | CM084926.1 | 3597 | 3326 |
| GEFB01005208 | Glycoprotein hormone (GPH) | 19 | 5112 | CM084939.1 | 32323 | 6379 |
| GEFB01030964 | Inotocin | 3 | 2465 | CM084934.1 | 4175 | 180 |
| None found | Insulin-like peptide (ILP) | – | – | – | – | – |
| GEFB01016835 | Leucokinin (LK) | – | – | – | – | – |
| GEFB01024737 | Myosuppressin (MS) | 2 | 1269 | CM084915.1 | 22663 | 218 |
| GEFB01022366 | Neuropeptide F (NPF) | 2 | 1299 | CM084913.1 | 29579 | 242 |
| GEFB01009443 | Pigment dispersing hormone (PDH) | 14 | 1743 | CM084933.1 | 119695 | 108 |
| GEFB01004771 | Proctolin (Proc) | 9 | 2007 | CM084926.1 | 48180 | 1615 |
| GEFB01015867 | Pyrokinin (PK) | 7 | 1278 | CM084919.1 | 13337 | 58 |
| GEFB01027769 | Red pigment concentrating hormone (RPCH) | 9 | 1590 | CM084920.1 | 175091 | 74 |
| GEFB01016897 | RYamide (RYa) | 3 | 1199 | CM084951.1 | 33842 | 91 |
| GEFB01013521 | Short neuropeptide F (sNPF) | – | – | – | – | – |
| GEFB01030224 | SIFamide (SIFa) | 7 | 1741 | CM084959.1 | 32378 | 161 |
| GEFB01033477 | Sulfakinin (SK) | 9 | 3936 | CM084943.1 | 47245 | 769 |
| GEFB01026365 | Tachykinin-related peptide (TRP) | 9 | 2455 | CM084942.1 | 22666 | 465 |
| GEFB01026709 | Trissin | 4 | 1491 | CM084935.1 | 36228 | 177 |

**Figure 4.**
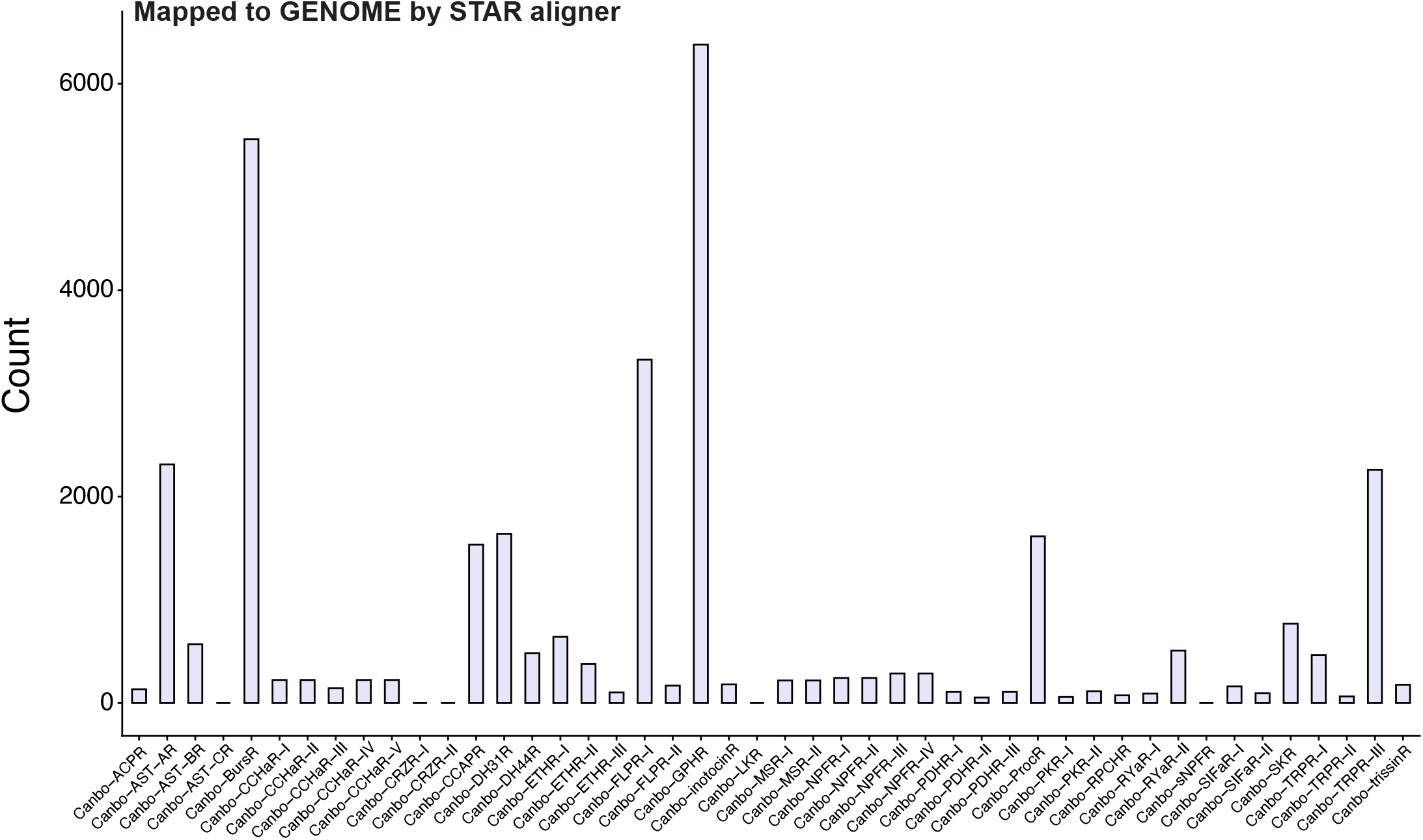
Transcript abundance of neuropeptide receptor genes in the *Cancer borealis* STG. Gene-level transcript abundance was quantified using the STAR aligner and featureCounts. Bars represent normalized transcript counts for selected neuropeptide receptor gene families involved in neuromodulatory signaling within the STG.

Most of these identified peptide receptors show some level of expression in the STG (Figure 4; 6). Well known peptide modulators of STG circuit function are represented among the more highly expressed transcripts including the putative allatastatin (AST), crustacean cardioactive peptide (CCAP), FLRFamide (FLPR), proctolin (ProcR), and tachykinin-related peptide (TRPR). Others that show fairly high abundance in STG that have been less well characterized include bursicon (BursR), diuretic hormone (DH31R), glycoprotein hormone (GPHR), and sulfakinin (SKR). It is worth noting that none of these receptor protein subtypes (Figure 4) have been functionally characterized in the STG, and these identifications are solely by sequence homology which is always a limitation for functional interpretation.

## Conclusions

By integrating chromosome-scale genomics with transcriptomic validation, this study reveals the structural and evolutionary foundations of neuronal gene diversity in *Cancer borealis*. Gene duplication, intronic expansion, and regulatory evolution emerge as key mechanisms enabling functional robustness and flexibility in crustacean neural circuits. These results establish a high-resolution genomic resource that bridges molecular genetics and systems neuroscience, supporting future studies of neural function, plasticity, and the evolution of nervous system complexity. RNAseq analysis of the *C. borealis* stomatogastric ganglion provides insight into which neuronal-related genes are expressed and to what relative abundance that may shed additional light on function in this well-studied model system.

## Data Availability

All RNA-seq data generated and analyzed in this study have been deposited in the NCBI Sequence Read Archive (SRA) BioProject ID PRJNA1505596. All scripts used for data processing and analysis are publicly available at https://github.com/murugesanraj/c-borealis-stg-genome-annotation.

## Acknowledgments

The authors acknowledge the Hellbender High Performance Computing facility at the University of Missouri for providing computational resources and technical support that made this work possible.

